# Within-host rates of insertion and deletion in the HIV-1 surface envelope glycoprotein

**DOI:** 10.1101/2023.09.19.558348

**Authors:** John Palmer, Vlad Novitsky, Roux-Cil Ferreira, Art F. Y. Poon

**Affiliations:** Department of Pathology and Laboratory Medicine, Western University, London, ON, Canada; Department of Immunology and Infectious Diseases, Harvard School of Public Health, MA, USA; Department of Microbiology and Immunology, Western University, London, ON, Canada; Department of Computer Science, Western University, London, ON, Canada

## Abstract

Under selection by neutralizing antibodies, the HIV-1 envelope glycoprotein gp120 undergoes rapid evolution within hosts, particularly in regions encoding the five variable loops (V1-V5). Indel polymorphisms are abundant in these loops, where they can facilitate immune escape by modifying the length, composition and glycosylation profile of these structures. Here, we present a comparative analysis of within-host indel rates and characteristics within the variable regions of gp120. We analyzed a total of 3,437 HIV-1 gp120 sequences sampled longitudinally from 29 different individuals using coalescent models in BEAST. Next, we used Historian to reconstruct ancestral sequences from the resulting tree samples, and fit a Poisson generalized linear model to the distribution of indel events to estimate their rates in the five variable loops. Overall, the mean insertion and deletion rates were 1.6 × 10*^−^*^3^ and 2.5 × 10*^−^*^3^/ nt / year, respectively, with significant variation among loops. Insertions and deletions also followed similar length distributions, except for significantly longer indels in V1 and V4 and shorter indels in V5. Insertions in V1, V2, and V4 tended to create new N-linked glycosylation sites significantly more often than expected by chance, which is consistent with positive selection to alter glycosylation patterns.

## Introduction

Human immunodeficiency virus type 1 (HIV-1) exhibits one of the fastest rates of molecular evolution measured in the natural world [1]. Key characteristics of HIV-1 that drive its rapid evolution include a high mutation rate associated with the error-prone reverse transcriptase encoded by the virus, a short generation time of roughly *∼* 2.5 days, and large population size with about 10^10^ to 10^12^ virions produced every day [2, 3]. HIV-1 evolution is shaped by different processes operating within and between hosts. Between-host evolution is predominantly shaped by stochastic processes associated with HIV-1 transmission [4]. For example, new infections are frequently established by a single copy of the virus, largely sampled at random from the previous host [5]. In contrast, selection within hosts drives the accumulation of mutations that enable the virus to escape the host-specific immune response. The neutralizing antibody response, for instance, can select for escape mutations associated with the surface-exposed portions of the HIV-1 envelope glycoprotein gp120 [2]. The gene encoding HIV-1 gp120 has five hypervariable regions (V1-V5) that yield unstructured (disordered) loops that can shield more conserved regions of the glycoprotein from neutralizing antibodies [6]. In addition, gp120 is heavily modified by N-linked glycosylation, the post-translational attachment of complex carbohydrates to asparagine residues, that results in a ‘glycan shield’ against neutralizating antibodies.

Mutations arise through multiple mechanisms, specifically nucleotide substitutions, insertions and deletions. When we observe homologous sequences of different lengths, it is not possible to determine whether an insertion or deletion has occurred without reconstructing the ancestral state. Hence, in the absence of additional information, insertions and deletions are conflated under the portmanteau ‘indels’. The rates and distribution of nucleotide substitutions in HIV-1 gp120 have been studied extensively [2, 7, 8]. However, the comparative methods used in these studies generally discount indels as missing data, *i.e.*, alignment gaps. Consequently the role of indels in virus adaptation, transmission and pathogenesis is poorly understood. The position and number of N-linked glycans in HIV-1 gp120, which tend to be associated with the hypervariable loop regions, are often modified through indel mutations that can have substantial effects on neutralization sensitivity [9–11].

In previous work [12], we reported measurements of indel rates in the variable loops of HIV-1 gp120 from a phylogenetic analysis of HIV-1 sequences from over 6,000 individuals. Here, we extend this work to quantify these indel rates within hosts through a time-scaled phylogenetic analysis of intra-patient HIV-1 sequence variation. We postulate that the observed indel rates will be higher within hosts because a greater number of individual mutations can be observed before they are removed by purifying selection [13]. In contrast, indels observed at the among-host level tend to be mutations that have increased to substantial frequencies in a given host. For example, we seldom observed indels at an among-host level that induced frame-shifts in HIV-1 gp120 [12]. Furthermore, we use recent advances in ancestral reconstruction methods to distinguish insertion from deletion events. We use these reconstruction results to characterize the composition of insertion and deletion sequences with respect to nucleotides and potential N-linked glycosylation sites (PNGSs).

## Methods

### Data collection

Our methodological workflow is summarized in Figure 1. First, we queried the Los Alamos National Laboratory (LANL) HIV sequence database (http://www.hiv.lanl.gov, accessed December 4, 2018) for HIV-1 gp120 sequences derived from longitudinally sampled patients. The resulting data comprised a total of 11,265 anonymized sequences representing 25 unique patients and 29 published studies. To maximize our ability to fit a molecular clock, we checked that each data set contained more than 100 sequences, spanned a time period greater than 200 days, and comprised five or more unique sampling time-points. We also confirmed that every sequence was annotated with a full sample collection date (year, month and day). Next, we referred to corresponding publications and GenBank entries to assess the nucleic acid type (DNA versus RNA) associated with each sequence. We excluded sequences derived from HIV-1 DNA as these may skew clock estimates, particularly when a patient is on suppressive ART. In addition, we incorporated an additional 2,541 HIV-1 RNA gp120 sequences collected from 42 patients in Botswana: 1,966 of these sequences representing 32 patients were published with a previous study [14], and the remaining 574 sequences were associated with 10 patients in another study [15]. These sequence data were generated using single genome amplification (SGA) and covered HIV-1 gp120 from regions V1 to C5 (HXB2 nucleotide coordinates 6615 to 7757). After initial screening, our resulting dataset contained 6,717 sequences stratified across 44 patients.

**Figure 1:**
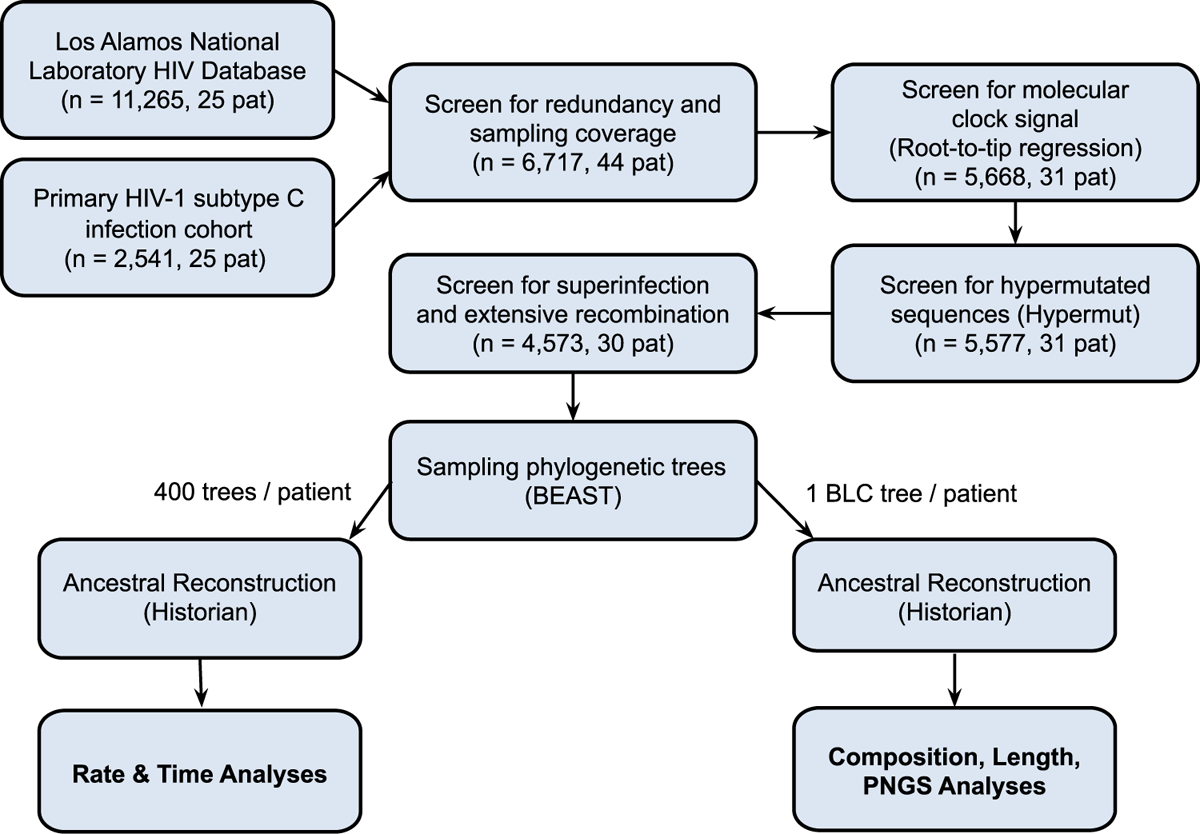
Summary of data sourcing, filtering, and analysis steps used in this study. The numbers of patients (pat) and sequences are updated at each major step. BLC = branch length consensus.

We performed pairwise alignment of each nucleotide sequence against the gp120 gene in the HXB2 reference sequence (Genbank accession K03455) to remove nucleotides outside this target region, using the Gotoh algorithm as modified by Altschul and Erickson [16] (http://github.com/ ArtPoon/gotoh2). Next, we removed gap characters from the trimmed sequences, translated them into amino acid sequences, and aligned the results to the HXB2 gp120 protein sequence. Based on gap positions in the resulting amino acid alignment, we inserted gap triplets into the appropriate locations of the patient’s nucleotide sequence in order to generate codon-aware nucleotide alignments. We recorded the start and end positions of the variable loops for each of these pairwise alignments. Finally, we used MAFFT (version 7.271) [17] to generate a multiple alignment of all trimmed sequences per individual.

### Data filtering

We used RAxML (version 8.2.11) [18] to reconstruct maximum likelihood phylogenies for each set of sequences per individual. To root the resulting phylogenetic trees, we used the root-to-tip regression function *rtt* in the R package *ape* [19]. In addition, we used the x-intercept of this regression to obtain a rough estimate of the date at the root of the tree, which we assumed corresponds to the start of infection. We evaluated support for a molecular clock for each dataset by calculating the *R*^2^ associated with the regression model and removed data sets with values below 0.3. Next, we screened for G to A hypermutation using an implementation of the Hypermut method [20] in Python. For each data set, we used the consensus of sequences from the first time point as a reference sequence to infer the directionality of nucleotide substitutions (*e.g.*, G to A). Finally, we manually examined the alignments and trees for evidence of co-infection or superinfection, based on the presence of distinct subpopulations separated by a deep branch. This step flagged one data set (LANL ID 111848), which we subsequently confirmed was sampled from an individual with a highly heterogeneous infection and extensive recombination [21], and was therefore excluded from further analysis. After these filtering steps, our data comprised 4,573 sequences from 30 individuals in total.

### Phylogenetic analysis

We used BEAST (version 1.10.4) [22] to sample trees from the posterior distribution. For each of the *n* = 30 individual data sets, we ran four replicate chain samples for 2 × 10^8^ iterations each. Two chains were run under a constant population size tree prior, and the other two were run with the Skygrid tree prior [23] that accommodates variation in population size (coalescence rates) over time. Based on the convergence characteristics of preliminary runs, we selected 20 grid points (population sizes) for the Skygrid model, and constrained the model to a maximum height of 125% of the difference between the estimated root date (from the preceding RTT analysis of the ML tree) and the most recent sample collection date. We used a Tamura-Nei (TN93) model of nucleotide substitution with a gamma distribution discretized into four rate categories to model rate variation among sites, and an uncorrelated lognormal (UCLN) relaxed clock model for rate variation among branches [24]. To improve convergence, we used previous estimates for HIV-1 gp120 [3] to calibrate the lognormal prior for the mean hyperparameter of the UCLN (*µ* = 1.2 × 10*^−^*^4^, *σ* = 1.5 × 10*^−^*^4^). Even so, one of the data sets did not converge for either model. We used Bayes factors to select between constant and Skygrid coalescent chain samples for each individual data set, except in 9 of 60 (15%) chain samples where the posterior trace was too irregular to have confidence in the associated model likelihoods. In these cases, we selected the model with better evidence of convergence between replicate chain samples. Overall, the Skygrid model was supported in 19 out of 29 data sets, while there was insufficient evidence to reject the constant population size model in remaining 10.

### Indel rate estimation

We downsampled trees from the chain samples and used Historian (version 0.1) [25] to reconstruct ancestral sequences along with the associated insertion and deletion events. For indel rate and timing analyses, we ran Historian on a random sample of 50 trees per chain sample, producing a total of 100 trees per patient (Figure 1). For the remaining analyses of nucleotide sequences (*i.e.*, lengths, compositions, PNGS interactions), we used samples of 100 trees per chain sample, or 200 trees per patient. Our fully finalized outputs from Historian analysis included 2,892 sequences stratified across 27 patients. When stratified by group M subtype, there were 2,585 subtype C (24 patients), 216 subtype B (2 patients) and 91 subtype A1 sequences (1 patient).

Next, we extracted insertions and deletions by recursively iterating through intra-patient trees to compare adjacent sequences on each branch, *i.e.* every ancestor-descendant pair. We specifically scanned the variable regions within the Historian output alignment using locations determined from our previous in-frame pairwise alignment with HXB2 gp120. Insertions presented as a string of one or more nucleotides only found in the descendant, while deletions were only present in the ancestor. To estimate insertion and deletion rates, we extracted the count and evolutionary time (time-scaled length) associated with every branch on 100 randomly sampled trees from each of 27 patients (*n* = 2,585 sequences total). We then fit a log-linked generalized linear model (GLM) to these quantities, modeling the number of insertions or deletions as a Poisson-distributed outcome *Y* with probability *P*(*Y|λ, t*) = (*λt*)*^Y^ e^−λt^/Y* !, where *λ* is the rate of insertion or deletion, and *t* is the branch length in units of time. To obtain confidence intervals, we replicated this regression analysis across ancestral reconstructions for all sampled trees. We also recorded the cumulative edge length from the tree root to the midpoint of every branch to quantify the estimated time since the start of infection for every inferred insertion or deletion event.

### Analysis of indel sequences

Given the reconstructed insertion and deletion sequences, we examined the nucleotide composition and length distribution of the indels, and their effects on potential N-linked glycosylation sites (PNGSs). First, we used a randomization test to evaluate whether the nucleotide composition of insertions or deletion sequences were significantly different in comparison to the flanking ‘non-indel’ sequence in the variable loop. For every indel, we randomly sampled 100 sub-strings of equivalent length from the inferred ancestral variable loop sequence to generate a null distribution of nucleotide frequencies within indels. We tested for significant differences using a randomization test which involved: (1) sampling each variable loop sequence 100 times for substrings matching the size of the indel; (2) pooling these substrings together; (3) calculating the nucleotide proportions on this null distribution of variable loop samples, and; (4) testing whether observed insertion/deletion nucleotide proportions were above or below the 95% quantiles of this distribution.

Next, we categorized insertions and deletions by length into nine categories, stratified by variable loop, and used the *mosaic* function in the R package *vcd* [26] to detect significant departures from the expected frequencies. This function computes the Pearson *χ*^2^ residuals in the contingency table of variable loops against length categories to detect counts that were significantly higher or lower than expected. Finally, we screened variable loop sequences for PNGSs using the regular expression ‘*N*[^*P*][*ST*][^*P*]’, where ‘[^*P*]‘ will match any amino acid except proline. We counted the number of PNGSs in ancestral sequences before and after every reconstructed indel event. We then compared the ‘observed’ changes in the number of PNGS induced by indels to a null distribution, which we generated by randomizing the placement of the same indels in their respective variable loop sequences for 500 replicates.

## Results

### Indel rate estimates

We generated posterior samples of phylogenies relating a total of 2,892 longitudinally-sampled HIV-1 gp120 sequences from 29 different individuals, and then reconstructed ancestral sequences of 100 random trees from each posterior sample per patient to extract insertions and deletions. We then fit a Poisson generalized linear model to time and indel count data extracted from these 2,700 trees (27 patients × 100 trees) to estimate the insertion and deletion rates in the five gp120 variable loops. The global mean insertion rates along terminal and internal branches were 1.6 × 10*^−^*^3^ and 2.5 × 10*^−^*^3^ events/nt/year, respectively (Figure 2). Across all variable loops and types of branches, insertion rates ranged between 2.9 × 10*^−^*^5^ (V3) and 5.4 × 10*^−^*^3^ events/nt/year (V1; Figure 2). The average deletion rate estimated along terminal branches was 3.2 × 10*^−^*^3^ events/nt/year, while the internal branch mean was 3.5 × 10*^−^*^3^ events/nt/year. Collectively, deletion rates ranged between 2.0 × 10*^−^*^5^ and 7.0 × 10*^−^*^3^ events/nt/year (Figure 2). Based on the separation of 95% confidence intervals, rates of both insertions and deletions were significantly highest in V1 and V5, and lowest in V3. Overall, indel rates were generally higher on internal branches versus terminal branches, consistent with the removal of mutations by purifying selection. For comparison, the overall mean nucleotide substitution rate from the same data was 2.2 × 10*^−^*^2^ events/nt/year with a range from 7.7 × 10*^−^*^3^ to 5.8 × 10*^−^*^2^ (Supplementary Figure S1).

**Figure 2:**
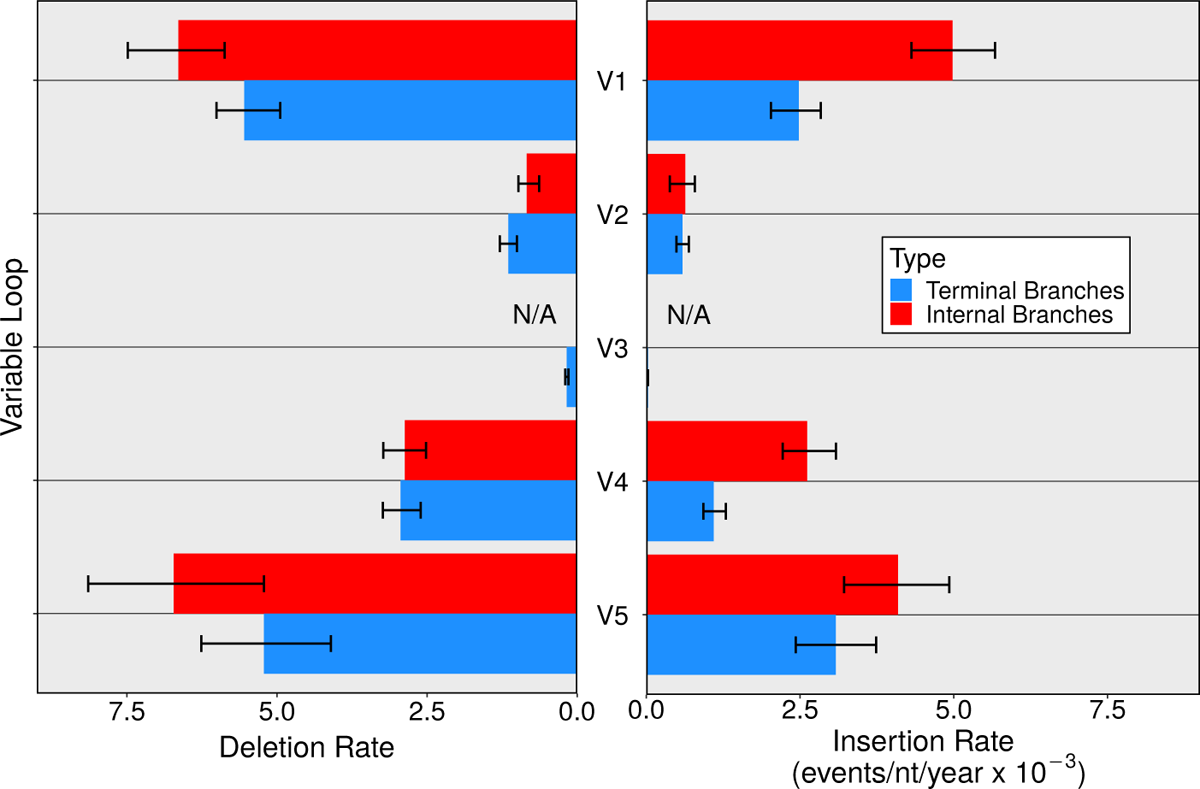
Mean within-host insertion and deletion rates in the five gp120 variable loops of HIV-1 subtype C. Each rate was estimated using a Poisson generalized linear model fit to time and indel count data extracted from 2400 phylogenetic trees (24 patients x 100 trees per patient). Rate are further stratified into those estimated along the terminal and internal branches within phylogenetic trees. Error bars represent the 95% confidence intervals generated using 100 bootstrap replicates.

### Indel timings

While extracting indels from within-host phylogenies, we also collected the cumulative time-scaled branch length from the tree root to the midpoint of the branch on which the indel occurred. We used these time values to approximate the distribution of insertion and deletion timings over the course of infection shown in Figure 3. Relatively few insertions and deletions accumulated in the first approximate 100 days of infection relative to the next 900 days (Figure 3). Following the first 100 days, insertions and deletions both appear to accumulate at a relatively consistent rates, although confidence intervals became progressively wider with decreasing sample sizes over time (Figure 3). Interestingly, deletions accumulated at a frequency of approximately 1.5 events/100 days / individual, while insertions accumulated at roughly 1.0 events/ 100 days / individual (Figure 3). Deletions were significantly more frequent than insertions during the first 600 days of infection. Based on the 95% confidence intervals, we estimated cumulative numbers of between 5 to 10 insertions and 9 to 18 deletions across the five gp120 variable loops within the first 1,000 days (*∼* 3 years) of infection.

**Figure 3:**
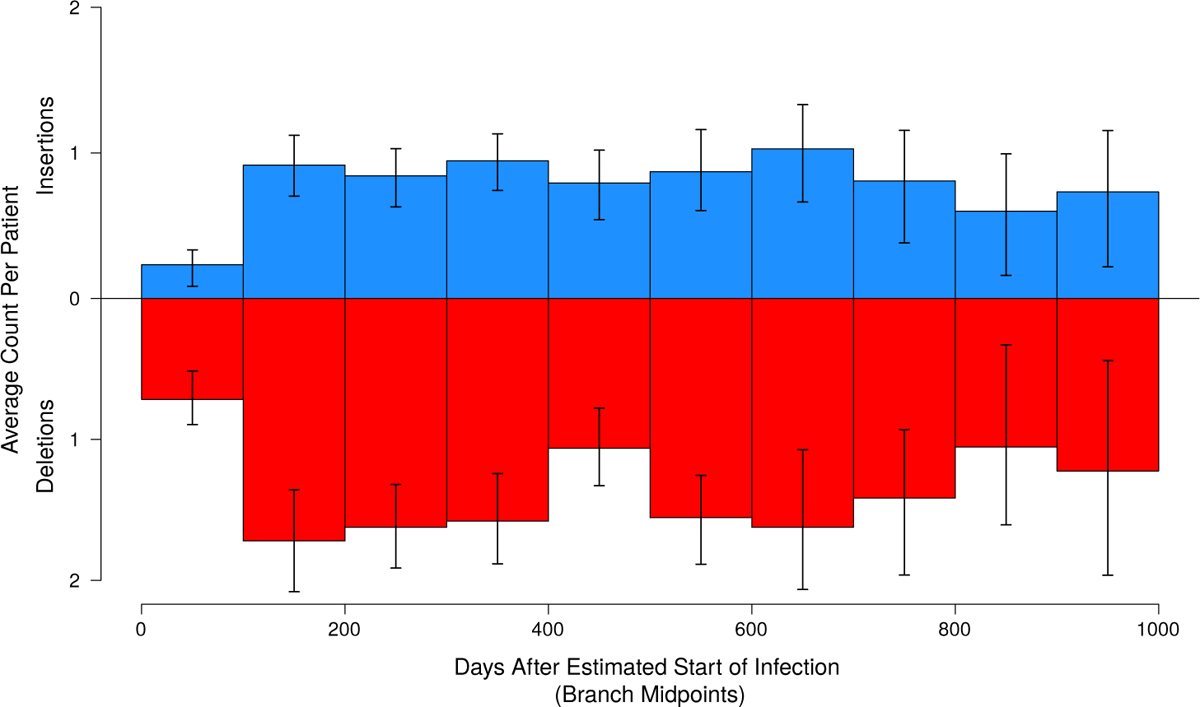
The mean counts of insertions (blue) and deletions (red) collected per patient every 100-day interval since the estimated start of infection (tree root). Error bars represent the 95% confidence intervals generated using 100 bootstrap replicates. Timings are measured by the cumulative time-scaled edge distance from the root of the tree to the midpoint of the branch where the indel occurred. To account for differences in maximum tree height, we applied a correction factor at higher time intervals as fewer datasets became available.

### Indel lengths

For remaining analyses of indel lengths, compositions, and PNGS interactions, we analyzed indels retrieved from a sample of 20 posterior trees per patient across HIV-1 subtype A1, B, and C. We first examined the average lengths of variable loop insertions and deletions as shown in Figure 4. Interestingly, there were significantly elevated counts of both insertions and deletions longer than 12 nt in V1, and exactly 12 nt long in V4. In contrast, V5 had significantly increased counts for both insertions and deletions of 3 or 6 nt, and a poor tolerance for indels longer than 12 nt (Figure 4). Counts of insertions or deletions in V3 with lengths longer than 3 nt were significantly lower than expected. Frameshift-inducing lengths were very rare in both insertions and deletions, comprising 3.9% and 3.7% of these populations, respectively (Figure 4).

**Figure 4:**
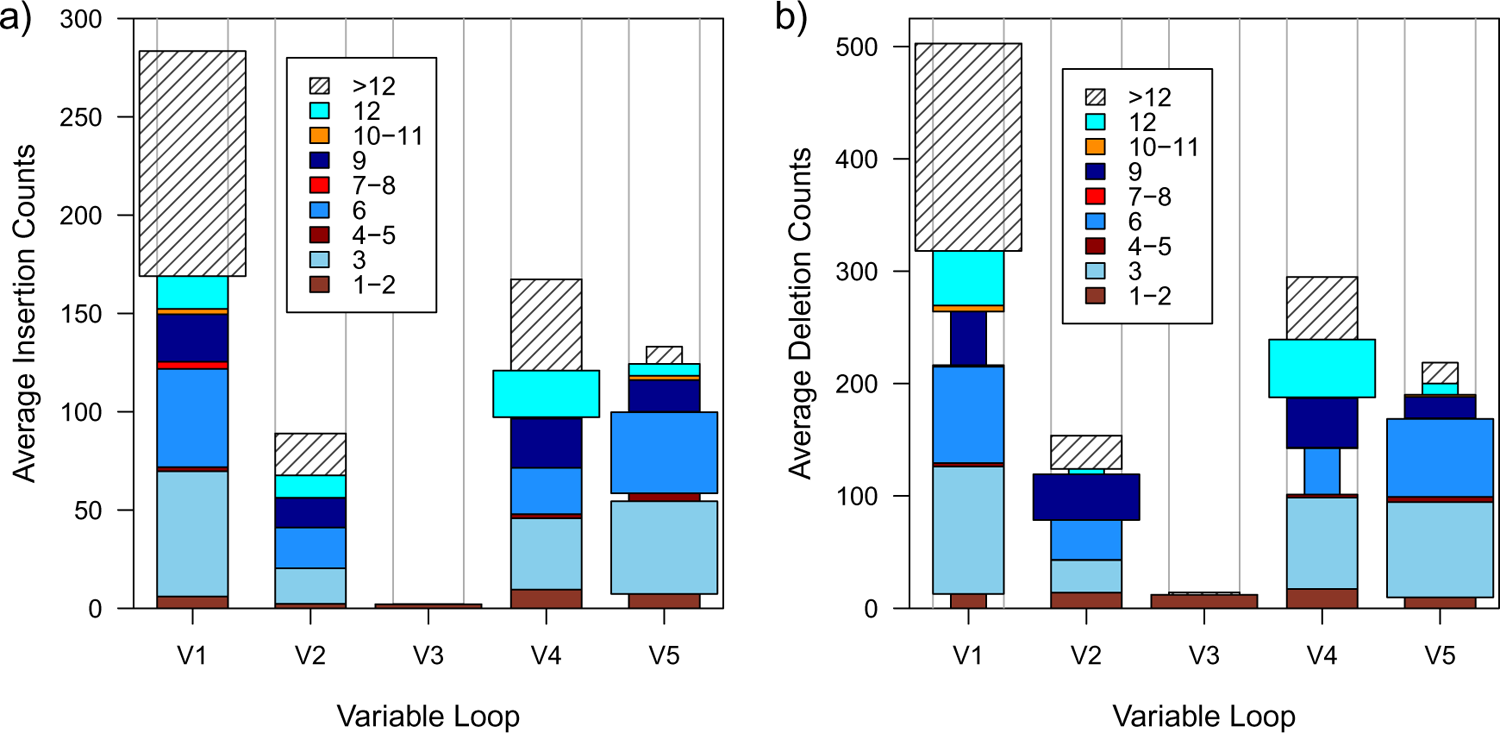
Mean counts of different insertion (a) and deletion (b) lengths recovered in the five variable loops of HIV-1 gp120. Block heights denote the mean number of indels in the given length category across 200 trees sampled per patient. Indel counts are stratified into nine length categories distinguished by colour or line shading (see inset legend). Pearson *χ*^2^ residuals (standardized differences between observed and expected counts) were generated for every length bin and used to detect significant differences. Bins that are wider and narrower that the outlined column margins indicate counts that are significantly higher and lower than expected, respectively, based on Pearson *χ*^2^ residuals (*α* = 0.05). Bins matching the column margins in width showed no significant difference.

### Indel compositions

Next, we sought to determine whether indel compositions differed significantly from those of their surrounding variable loop regions. We investigated these compositional differences by estimating nucleotide proportions within insertions and deletions, and compared them to the flanking regions in the inferred ancestral sequences on branches associated with the indel events. Overall, average nucleotide compositions of both insertions and deletions showed little deviation from those of their variable loops of origin, as shown in Figure 5. Similar results were observed when investigating indel dinucleotide proportions, which also showed high similarity to those within the variable loops (Supplementary Figures S4 and S5). Indels and variable loops both demonstrated an intolerance for CpG dinucleotides as expected, given their role in sequence methylation. Relative to surrounding non-indel sequences, there were slightly elevated proportions of guanine in insertion sequences, and slightly decreased proportions of thymine in both insertions and deletions. However, a two-sided randomization test with *α* = 0.05 found neither of these deviations to be significant (Figure 5). Further stratification of nucleotide proportions by variable loop revealed that these higher proportions of G in insertions are predominantly found in V2 and V5 (Supplementary Figure S2a). Guanine proportions in insertions were relatively higher than those in deletions, though we did not investigate this further given the small magnitude of this difference (Figure 5).

**Figure 5:**
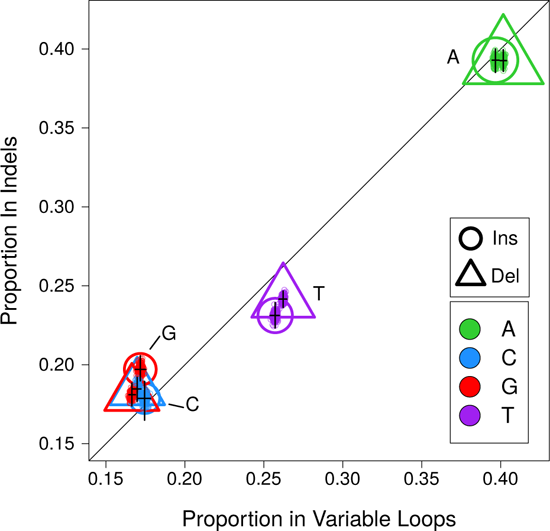
Nucleotide compositions of variable loop insertions (circles) and deletions (triangles) relative to their surrounding non-indel variable loop sequence. Each shape represents the mean proportions of a nucleotide within indels (*y*-axis) or the remainder of the variable loop (*x*-axis). Shape area is scaled in proportion to the overall nucleotide frequency. Error bars represent the 95% confidence intervals for nucleotide proportion estimates based on 100 bootstrap replicates. Deviations from the center line indicate differences between the nucleotide proportions of indels relative to their surrounding variable loop sequences.

### Indel-induced PNGS changes

Finally, we investigated whether indels add or remove potential N-linked glycosylation sites (PNG-Ss) more often than expected by chance. We located PNGSs within the variable loops of 20 posterior trees using regular expressions and observed how PNGS numbers were altered by indels. On average, insertions in V1, V2 and V4 increased PNGS counts significantly more often than what would be expected were they to be placed randomly. Specifically, the reconstructed insertions in V1, V2, and V4 resulted in the creation of 0.42, 0.2 and 0.29 PNGSs on average. In contrast, randomized placement of the same insertion sequences into the ancestral V1, V2 and V4 loops resulted in a net loss of 0.32, 0.23 and 0.53 PNGSs on average (Figure 6). Deletions in these variable loops also tended to disrupt PNGSs, though these changes were relatively comparable to those expected from random placement (Figure 6). The reconstructed insertions in V3 and V5, and deletions in V2, V3, and V5, did not induce a net change in the number of PNGSs that was measureably different from what we expected from randomization. In particular, none of the reconstructed insertions in V3 affected the small number of PNGSs in this variable loop.

**Figure 6:**
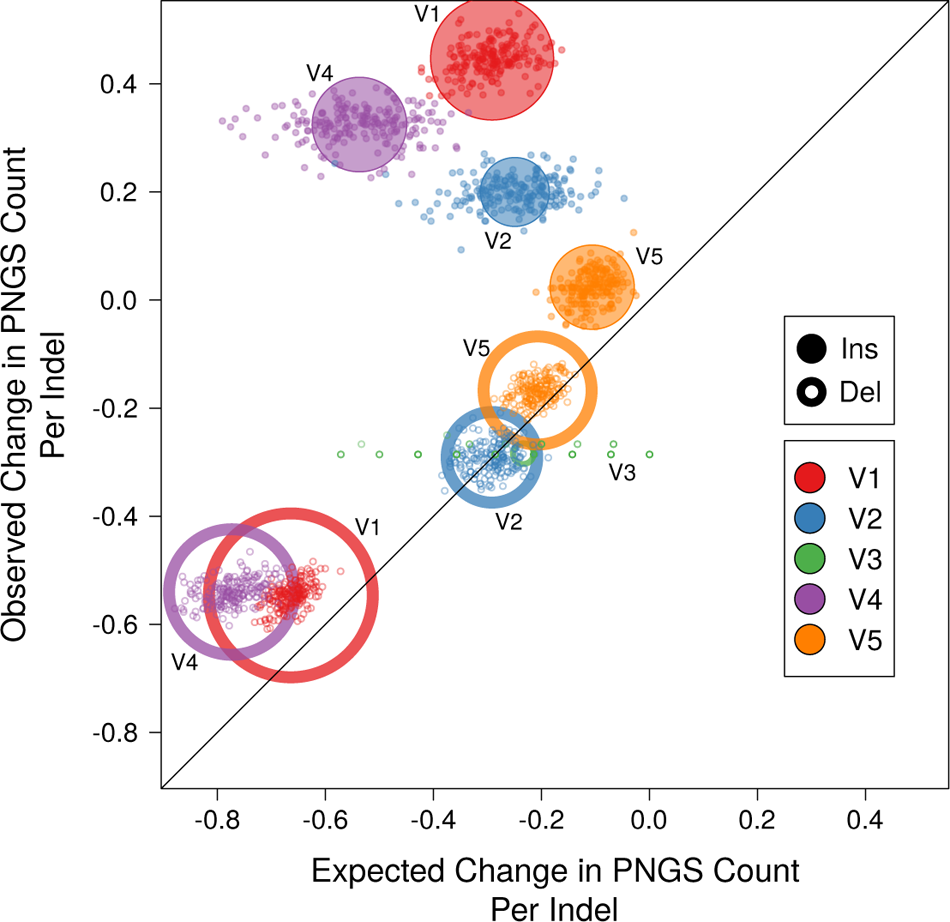
Indel-induced changes to counts of potential N-linked glycosylation sites (PNGSs) in the variable loops. Each point represents the mean change in the number of PNGSs induced by ‘observed’ indel events in the data (*y*-axis) or randomized placement of indels (*x*-axis) for one of 200 bootstrap samples. Open and closed circles indicate the overall mean per event type (deletions and insertions, respectively) and by variable loop (indicated by colour). Circle areas are scaled in proportion to the number of indel events.

## Discussion

In this study, we provide the first estimates of indel rates in HIV-1 gp120 measured on a within-host scale. While Mansky and Temin [27] has estimated the rate of indels generated *de novo* across the HIV-1 genome in addition to nucleotide substitutions, the rates of indel evolution in patient-derived HIV-1 sequences have seldom been investigated. Here, we expand on our past work [12] by measuring the within-host indel rates for the gp120 variable loops of HIV-1 subtype C. We found that the variable loops have collective mean insertion and deletion rates of 1.96 × 10*^−^*^3^ and 3.21 × 10^3^ events/nt/year, respectively. Comparing these within-host indel rates (combined for both insertions and deletions) to our previous rate estimates among hosts [12], we observed that the within-host rates were significantly higher in V1, V2, V4, and V5, based on the 95% confidence intervals (Figure 2). In concordance with our expectations, our findings of higher indel rates within hosts than among hosts recapitulate the evolutionary trends observed in nucleotide substitution rates [3]. Interestingly, within-host indel rates exhibit a similar pattern of variation among variable loops relative to what we observed among hosts [12] (Figure 2), indicating that differences in the amount of purifying selection between these levels of evolution did not overwhelm variation among loops.

Higher deletion rates (Figure 2) along with similar length distributions between insertions and deletions (Figure 4) have interesting implications for the evolution of HIV-1 within hosts. Together, these findings suggest that the variable loops undergo a net reduction in the number of nucleotides over the course of infection, which is not consistent with previous reports of increasing lengths of variable loops from Derdeyn et al. [10] and Sagar et al. [28]. A potential explanation for this trend is the selection of shorter variable loops to enhance the efficiency of HIV-1 binding to susceptible immune cells during early stages of infection [29]. Our findings of roughly 1.9 times as many deletions as insertions observed in the first 400 days of infection offer additional support for this notion (Figure 3). Interestingly, Cheynier et al. [30] reported collecting four times as many deletions than insertions in V1 of a strain of SIV, which also suggests a significant presence of deletions at some point during infection. In light of our findings and previous ones, we speculate that HIV-1 infection sees an early, short-lived spike in deletions (loop shrinkage) followed by a gradual shift towards higher insertion prevalence (lengthening) later in infection as shown in past findings from Derdeyn et al. [10] and Sagar et al. [28].

Furthermore, we found that most variable loops had comparable terminal and internal branch rate estimates based on their 95% confidence intervals. Contrary to our expectations however, we found cases — namely, insertion rates in V1 and V4, and deletion rates in V2 — where rate estimates on internal branches were higher than those estimated on terminal branches. This could be caused by rapid, exponential population growth during early infection, which may produce shorter internal tree branches that elevate rate estimates in these regions [31]. In fact, we found groups of short internal branch in our posterior trees, indicating this as a likely driving force for higher internal branch rates. As discussed previously with deletions, this could also be explained by selective pressures driving the preferential accumulation of indels during early infection, potentially as a means to alter binding efficiency [29].

By investigating the distribution of indel timings, we also found a subtle, yet interesting, trend in variable loop indel timings during the first 1000 days of infection (Figure 3). After low counts in the first 100 days, indel counts climbed to higher values then appeared to remain consistent over the course of infection (Figure 3). Although results suggest a slight decline in indel counts toward the 1000 day mark, the high levels of uncertainty associated with results in this time period does not provide sufficient evidence to make concrete conclusions.

Unexpectedly, there were very few cases of frameshift-inducing insertions (3.9%) and deletions (3.7%), which suggest that our within-host observations are still subject to substantial purifying selection (Figure 4). Variable loop 3 demonstrated a complete intolerance for indels longer than 3 nt; however, this result is not surprising given its functional involvement in coreceptor binding and corresponding conserved nature [32]. Our findings of abundant long V1 insertions (*≥* 12 nt) align with previous findings that the V1V2 loop lengthens and gains PNGS over the course of infection in response to neutralizing antibody responses (Figure 4) [10, 28, 33]. It is interesting to note that insertions of this size (*≥* 12 nt) are capable of independently adding full PNGSs to the variable loop sequences (Figure 4) [9, 10, 28]. However, we also find high numbers of large deletions in V1, demonstrating that this loop has a high tolerance for both insertions and deletions and likely undergoes substantial changes in length. It is also surprising that we do not observed the same abundance of long indels in V2 considering: it is the longest of the five loops (120 nt), it exhibits similar characteristics to V1, and the previous reports of this loop accumulating numerous indels throughout infection (Figure 4) [2, 9, 28, 34].

We also found that the nucleotide compositions of both insertions and deletions were highly similar to those found in their variable loops of origin (Figure 5). These findings loosely suggest that the nucleotides comprising insertions originate from somewhere in the HIV-1 genome, given the distinctly high proportions of A and low proportions of C found in both sources (Figure 5) [35]. Regarding deletions, these findings suggest that there are no relative differences in the fitness costs of maintaining each of four nucleotides in the variable loop sequences, since all nucleotides have a roughly equal propensity to be deleted (Figure 5).

We found that insertions in V1, V2 and V4 tended to create PNGSs in these loops more frequently than expected by chance (Figure 6). In these loops, these stark differences between expected PNGS losses and observed gains suggest that insertions are under significant selective pressure to avoid disrupting existing PNGSs and plausibly add new ones (Figure 6). These findings align with previous reports that indels in V1, V2 and V4 tend to add PNGS as a likely means to escape neutralizing antibody responses over the course of infection [10, 28, 33, 36]. When taken in context of our findings of abundant long insertions in V1 and V4 (Figure 4), these results suggest that PNGS changes in these two loops are driven by long insertions (*≥* 12 nt), which have potential to contain entire PNGSs [10, 28, 36]. While deletions in V1 and V4 clearly tend to remove PNGSs, there is little evidence of selective pressures modulating their placement, given that their observed outcomes are highly similar to those expected from stochastic processes (Figure 6).

An important limitation to consider is our constraints placed on the prior distributions of hyperparameters describing the uncorrelated relaxed molecular clock model within the BEAST phylogenetic reconstruction. We narrowed these prior distributions due to the sampling of erroneously high values during BEAST MCMC, which significantly delayed analysis convergence. While introducing these constraints was necessary to achieve meaningful results, it is important to consider that they narrow our search of the parameter space. Posterior clock rate estimates may therefore, be biased towards our selected distribution, leading to an underestimation of their true variance. Lessened variation among clock rates may, in turn, reduce variation among time-scaled branch lengths in posterior trees, leading to an underestimation of variance associated with indel rate estimates.

Here, we perform a quantified investigation of indels in the gp120 variable loops on a within-host scale, which comprise an important evolutionary mechanism driving HIV-1 adaptation and immune escape. Indel rate estimates were higher on a within-host scale than those previously estimated among hosts, reflecting the same pattern found in nucleotide substitutions (Figure 2) [3, 12]. Interestingly, we found that frameshift-inducing insertions and deletions (3.9%, 3.7% respectively) were rare, demonstrating that purifying selection remained highly prevalent in our intrapatient datasets (Figure 4). Nucleotide compositions of insertions and deletions were essentially indistinguishable from those recorded in the surrounding variable loop sequences, loosely suggesting that insertions do not have a distinctly different origin and that nucleotides are not being selectively deleted (Figure 5). Importantly, we also find that insertions in V1, V2, and V4 are under selective pressure to introduce new PNGS sites, while deletions tend to remove PNGS in a roughly stochastic manner (Figure 6).

By estimating indel rates, we provide the first quantified measure of the within-host evolution introduced by indels in the gp120 variable loops. Given the suggested role of variable loop indels in immune escape, indel rates may correlate with HIV-1 disease progression or CD4^+^ cell decline, in a manner similar to substitution rates [37–39]. However, future research on indel rates will be needed to investigate these putative correlations. We hope that our quantification of variable loop indel rates will inspire future research into HIV-1 indel evolution and facilitate the incorporation of indels into evolutionary analyses.

## Acknowledgements

This work was supported in part by grants from the Natural Sciences and Engineering Research Council of Canada (NSERC RGPIN 05516-2018) and the Canadian Institutes of Health Research (CIHR PJT-155990). JP was supported by a Canada Graduate Scholarship from CIHR. RCF was supported by a Postdoctoral Fellowship in Genome Data Science from Ontario Genomics and the Canadian Statistical Sciences Institute.

