## Supplemental Figures for "Within-host rates of insertion and deletion in the HIV-1 surface envelope glycoprotein"

### Rates and Patterns of Indels in HIV-1 gp120 Within Hosts : Supplementary Material

#### Supplementary Figures

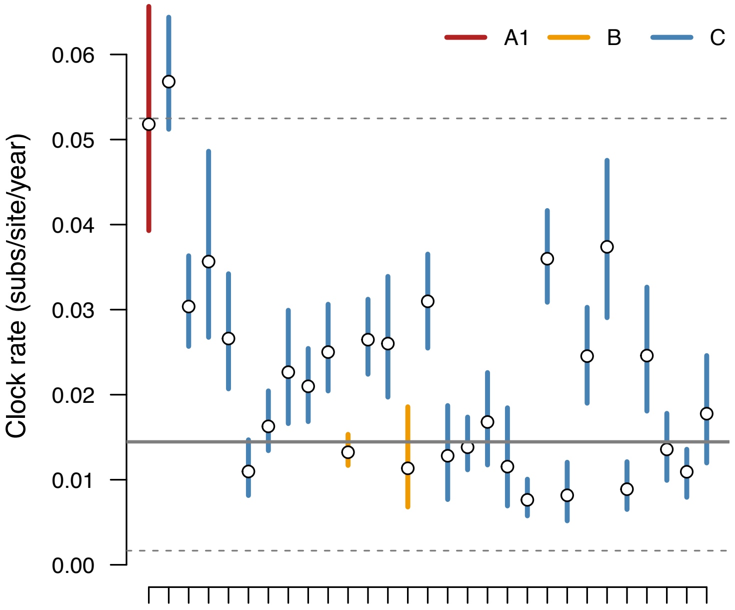


Figure S1: Estimates of molecular clock rates from BEAST analyses. Each point represents the mean estimate of the nucleotide substitution rate (molecular clock) from combining two BEAST chain samples for each individual. The samples were generated under a Skygrid prior for variation in coalescence rates (effective population size). Line segments represent the 95% quantile for clock rates, coloured by HIV-1 subtype (see inset legend). The solid and dashed horizontal lines represent the mean and 95% confidence interval, respectively, for estimates of the within-host clock for HIV-1 *env* from Alizon and Fraser (2013).

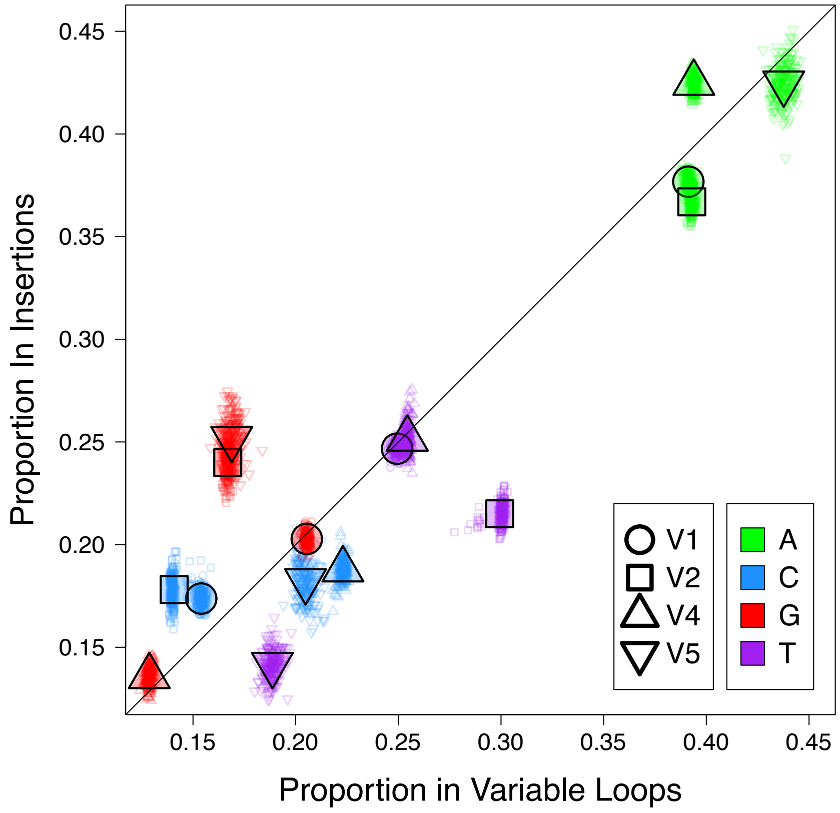


Figure S2: Nucleotide composition of insertion sequences relative to that of the surrounding variable loop sequence. Proportions of four nucleotides are further stratified by four variable loops: V1, V2, V4, and V5. Colors denote different nucleotides while unique shapes correspond to different variable loops. Proportions in V3 were excluded as extremely few insertion events were recovered in this loop. Deviations from the line of slope 1 denote differences in nucleotide proportions between insertions and that of surrounding variable loop sequences.

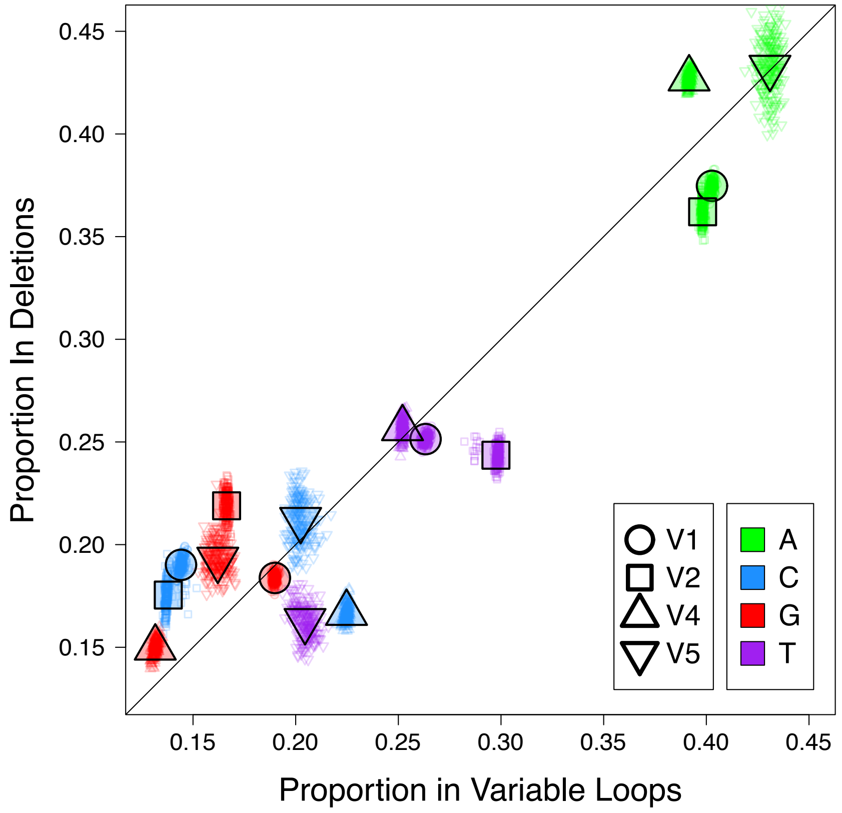


Figure S3: Nucleotide composition of insertion sequences relative to that of the surrounding variable loop sequence. Proportions of four nucleotides are further stratified by four variable loops: V1, V2, V4, and V5. Colors denote different nucleotides while unique shapes correspond to different variable loops. Proportions in V3 were excluded as extremely few insertion events were recovered in this loop. Deviations from the line of slope 1 denote differences in nucleotide proportions between insertions and that of surrounding variable loop sequences.

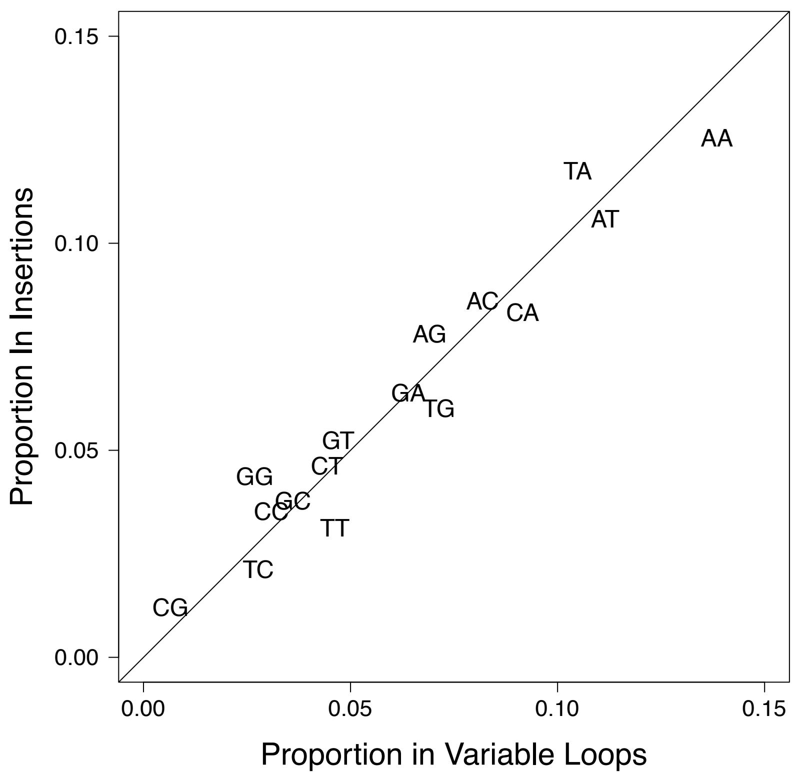


Figure S4: Dinucleotide proportions observed within insertion sequences (y-axis) relative to those proportions observed within their variable loop sequences of origin (x-axis). Deviations from the line of slope 1 denotes dinucleotide proportions that differed between recovered insertion sequences and their variable loops.

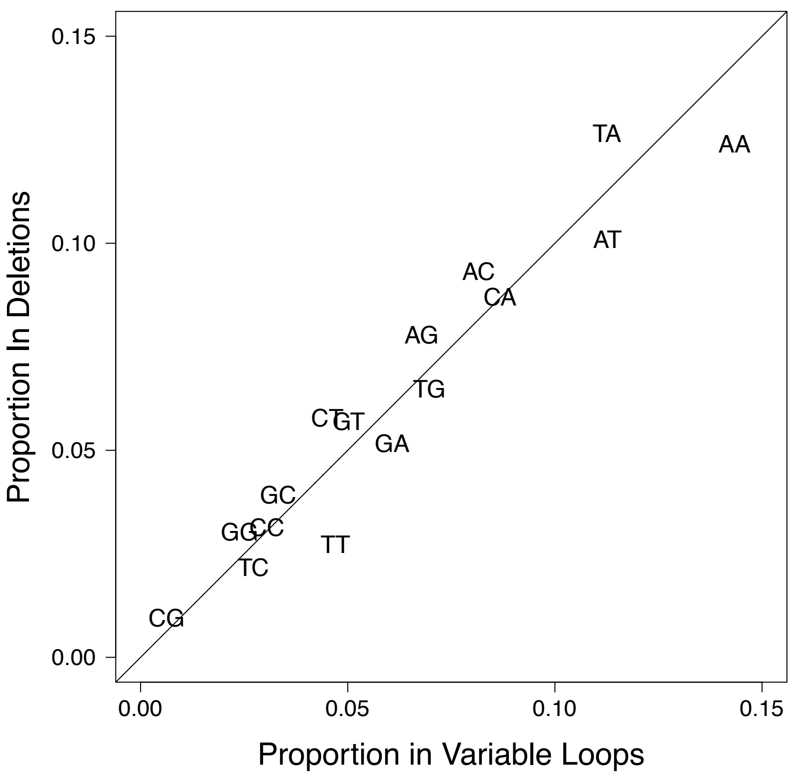


Figure S5: Dinucleotide proportions observed within deletion sequences (y-axis) relative to those proportions observed within their variable loop sequences of origin (x-axis). Deviations from the line of slope 1 denotes dinucleotide proportions that differed between deletion sequences and variable loops.
